# Cerebrospinal fluid (CSF) boosts metabolism and virulence expression factors in *Acinetobacter baumannii*

**DOI:** 10.1101/2020.07.13.201848

**Authors:** Jasmine Martinez, Chelsea Razo-Gutierrez, Casin Le, Robert Courville, Camila Pimentel, Christine Liu, Sammie E. Fuang, Alejandro J. Vila, Parvin Shahrestani, Veronica Jimenez, Marcelo E. Tolmasky, Scott A. Becka, Krisztina M. Papp-Wallace, Robert A. Bonomo, Alfonso Soler-Bistue, Rodrigo Sieira, Maria Soledad Ramirez

## Abstract

In a recent report by the Centers for Disease Control and Prevention (CDC), multidrug resistant (MDR) *Acinetobacter baumannii* is a pathogen described as an “urgent threat”. Infection with this bacterium manifests as different diseases such as community and nosocomial pneumonia, bloodstream infections, endocarditis, urinary tract, wound infections, burn infections, skin and soft tissue infections, and meningitis. In particular, nosocomial meningitis, a common complication of neurosurgery caused by extensively-drug resistant (XDR) *A. baumannii*, is extremely challenging to manage. Therefore, it is necessary to identify signals, such as exposure to cerebrospinal fluid (CSF), that trigger expression of virulence factors that are associated with the successful establishment and progress of this infection. While a hypervirulent *A. baumannii* strain did not show changes in its transcriptome when incubated in the presence of CSF, a low-virulence isolate showed significant differences in gene expression and phenotypic traits. Exposure to 4% CSF caused increased expression of virulence factors such as fimbriae, pilins, and iron chelators, and virulence as determined in various model systems. Furthermore, although CSF’s presence did not enhance bacterial growth, it was associated with an increase of expression of genes encoding transcription, translation, and the ATP synthesis machinery. Experiments to identify the active CSF component pointed to human serum albumin (HSA).

**Importance:** *Acinetobacter baumannii*, notorious for its multidrug resistant phenotype, overcomes nutrient deprived and desiccated conditions through its metabolic flexibility, pathogenic and physiological adaptability. Although this pathogen is commonly associated with respiratory infections, there have been a considerable amount of cases of *A. baumannii* bacterial meningitis. These infections are usually post-neurological surgery complications associated with high mortality rates ranging from 40 to 70%. This work describes interactions that may occur during *A. baumannii* infection of human cerebrospinal fluid (CSF). *A. baumannii’s* displays capabilities to persist and thrive in a nutrient-limited environment, which also triggers the expression of virulence factors. This work also further explores *A. baumannii’s* utilization of an essential component within CSF to trigger enhanced expression of genes associated with its pathoadaptibility in this environment.

## INTRODUCTION

*Acinetobacter baumannii* has emerged as an important pathogen due to its ability to resist multiple antibiotics, persist in hospital settings, and cause a wide variety of infections such as pneumonia, bacteremia, urinary tract infections, skin and soft-tissue infections, and meningitis (1–4). The acquisition of resistance to carbapenems by certain strains (carbapenem-resistant *Acinetobacter baumannii*, CRAB) increased the problematic nature of this pathogen (5), which has been qualified as an “urgent” threat in a recent report by the Centers for Disease Control and Prevention (6).

Bacterial meningitis, which is considered a medical emergency, is a serious infection that can cause permanent disabilities (brain damage, hearing loss, and learning problems) or death if untreated (7–9). Post neurosurgical *A. baumannii* meningitis can cause death or leave permanent sequelae and is usually associated with high mortality rates reaching up to 40 to 70 %. (10, 11). Illustrating the dangerous nature of these infections is the recent case of the *A. baumannii* infection of a 39-year-old man treated with external ventricular drainage of cerebrospinal fluid (CSF). Although the strain was susceptible to colistin at the time of detection, it quickly acquired resistance without losing virulence (12). This genetic plasticity, a consequence of its ability to acquire and integrate foreign DNA, gives *A. baumannii’s* a tremendous metabolic versatility that permits the bacterium to adapt and persist in harsh conditions (2, 13–17). *A. baumannii’s* success in causing numerous infections, where it gets in contact with different body components and fluids, must be the result of its capabilities to not only capture adequate genetic determinants but also regulate expression of the proper cell components (2, 13–15, 18–20). We have also previously documented that human serum albumin (HSA) and pleural fluid (HSA-containing fluid), affected *A. baumannii* behavior, triggering an adaptive response that modulates DNA uptake, cytotoxicity, immune evasion, stress responses and metabolism (18, 20–22).

Understanding the virulence of this bacterium requires a thorough comprehension of the general genotypic and phenotypic responses when it is exposed to the different body fluids. As part of our studies of the *A. baumannii* pathogenicity in relation to meningitis, we identified gene expression modifications when this bacterium is in contact with CSF impacting the phenotypic behavior of this organism.

## RESULTS AND DISCUSSION

### CSF treatment enhances the expression of genes involved in transcription and translation machineries, ATP production and specific metabolic pathways in *A. baumannii*

To further characterize *A. baumannii’s* transcriptomic response to human fluids, two different *A. baumannii* strains, A118 (low pathogenicity and high antibiotic susceptibility) and AB5075 (hypervirulent and multi-drug resistant), were exposed to CSF. Transcriptomic analysis of *A. baumannii* A118, using a fold-change cutoff of log_2_ > 1 (with adjusted *P*-value < 0.05), showed 275 differentially expressed-genes (DEGs), 7.76% of the total genes in the *A. baumannii* A118 reference genome. However, statistically significant changes were not observed when *A. baumannii* AB5075 was exposed to CSF. As, AB5075 is a hypervirulent and highly resistant, the lack of necessity to increase its pathogenicity can explain the absence of changes at the transcriptomic level in the tested conditions. Previous observations have shown that *A. baumannii’s* response to different stimuli is dependent on each particular strain’s degree of pathogenicity. Less pathogenic strains induced more changes in their phenotypic behavior to overcome the stressful environment and persist (23).

The analysis of *A. baumannii* A118 DEGs revealed an increase in the expression of many genes involved in the manifestation of genetic information and energy production machineries (Table S1). Notably, a large proportion of ribosomal protein genes are overexpressed upon exposure to CSF. Among the ribosomal protein associated genes, 47 out of 55 displayed a significant increase of expression of 2-fold or more. Coincidently, key translation genes such as those encoding elongation factors (EF) EF-G, EF-F and EF-P were also up-regulated. Concurrently, the main genes of the transcriptional machinery (RNA polymerase) were similarly overexpressed. The *rpoB and rpoC* genes, which code for the beta and beta’ subunits of RNA polymerase (core of the transcription machinery), were overexpressed with a log2fold just below 1. However, the gene encoding the alpha subunit was also up-regulated with a log_2_fold change of 1.48 (Table S1). In addition, genes important for energy production in the cell were also induced upon CSF exposure. The *atpIBEFHAGCD* locus, an operon encoding the FoF_1_-ATP synthase (the main ATP generator in the bacterial cell) displayed a ~3-fold transcriptional increase in expression (Table S1). These transcriptional responses suggest that *A. baumannii* responds to CSF by overexpressing machinery involved in gene expression (transcription and translation genes) and the main ATP generator, FoF1-ATP synthase. Also, CSF exposure induces the transcription of specific metabolic routes in *A. baumannii*. In particular, several dehydrogenases of tricarboxylic acid cycle (TCA) intermediates as well as a citrate symporter are significantly overexpressed, together with two proline symporters (Table S1).

Gene ontology (GO) was performed to identify molecular functions and biological pathways associated to *A*. *baumannii’*s adaptive responses to CSF. Consistently with the above mentioned DEGs, GO enrichment analysis revealed a statistically significant overrepresentation of the GO categories ATP synthesis coupled proton transport, translation, tricarboxylic acid cycle, and aerobic respiration by 13.8-, 8.8-, 4.9- and 4.3-fold, respectively (P-value < 0.05). Other authors found that exposure of *A. baumannii* strains to amikacin, imipenem, and meropenem was associated with an increase in the expression of genes involved in the TCA cycle, biosynthesis of amino acids, purines, and pyrimidines, as well as, the operons involved with ATP, RNA, and protein synthesis (24). Taken together, these results suggest that various stressful environments (nutrients availability and antibiotic treatment) induce expression of genes related to energy production, protein synthesis, and metabolism in *A. baumannii*. As a consequence of these transcriptomic changes, the bacterial cell must adapt to be able to survive and even thrive in these hostile conditions.

### CSF boosts specific metabolic routes without increasing growth in optimal growth conditions

The CSF-mediated upregulation of genes coding for the elements necessary for transcription, translation, expression and ATP synthesis was not accompanied by a decrease in generation time (Figure 1). These results suggest that cells respond to the presence of CSF by enhancing the expression of pathways that produce specific effects rather than increasing growth capacity. To better understand this behavior, we assessed the effect of CSF on cells growing in the presence of different carbon sources. *A. baumannii* A118 and AB5075 were cultured in minimal medium containing proline, glutamine, glucose, or citrate, all of which are utilized through the tricarboxylic acid cycle and whose expression was modified in the presence of CSF. Addition of CSF to a glutamine-containing minimal medium resulted in no change (*A. baumannii* A118) or a reduction (*A. baumannii* AB5075) of the growth rate (Figure S1). On the other hand, when the carbon source was proline, both strains grew at a higher rate (Figure S1). The transcriptomic data showed that type I glutamine synthetase (AbA118F_3228) and the proline symporter *putP* genes are upregulated by a log_2_fold change of 0.58 and 0.87 respectively. Studies on *Salmonella typhimurium* showed that *putP* codes for a proline permease, an integral membrane protein, that is the primary transport protein when this amino acid is the only carbon or nitrogen source (25). The CSF-mediated increase in abundance of the proline permease in *A. baumannii* may explain the increase in growth rate.

**Figure 1.**
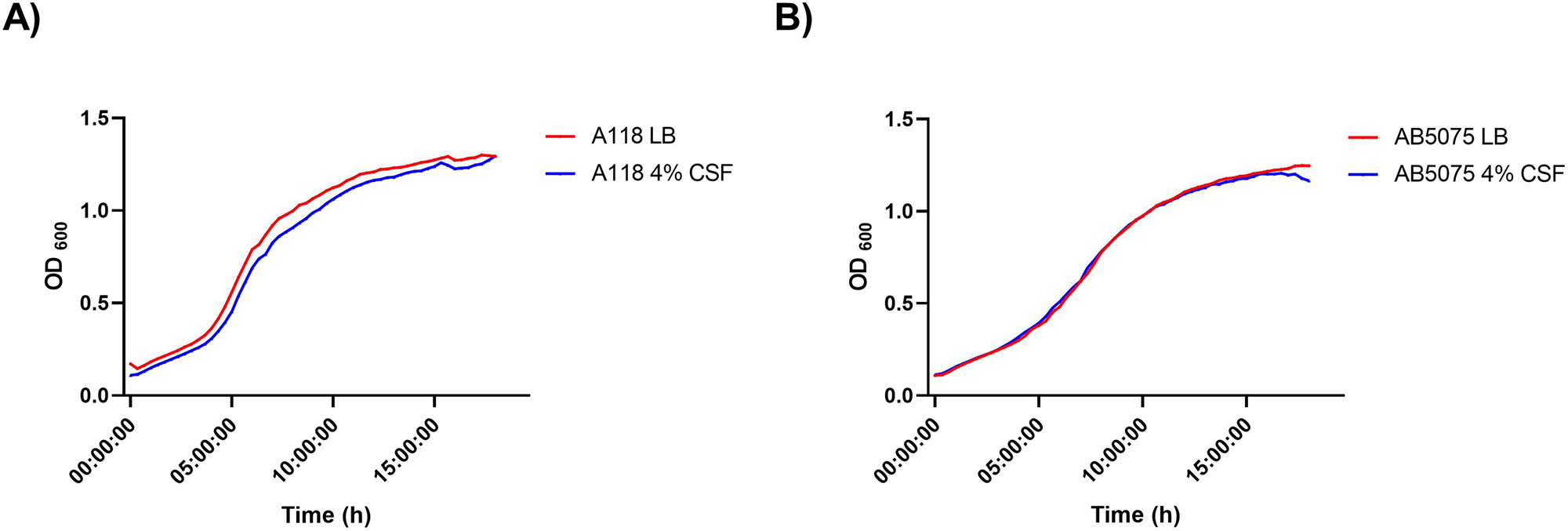
A118 and AB5075 growth curves in LB or LB plus 4% CSF. Strains A) A118 and B) AB5075 were grown in LB or LB supplemented with 4% CSF. Growth curves were conducted in triplicate.

CSF also induced an increase in the growth rate of both strains, although more pronounced in the case of *A. baumannii* AB5075, when cultured using citrate as sole carbon source (Figure S2). The transcriptomic analysis showed upregulation (log2fold change 0.62) of AbA118F_2664, a gene that encodes a CitMHS citrate-magnesium hydrogen complex symporter (Table S1). Proteins of the CitMHS family transport citrate-Mg^2+^ complex symport with one proton per complex molecule (26). It is of interest that increased citrate levels help survival of *A. baumannii* in certain conditions (22). Thus, the net effect of CSF might be an increase in expression of this transporter, which leads to higher citrate intracellular concentrations that may help by increasing the rate of growth. These results are consistent with previous observations that the presence of HSA, a major component of CSF, in the culture medium is correlated with elevated expression of the citrate symporter in *A. baumannii* (22). Further supporting this data, exposure of *A. baumannii* to pleural fluid (PF), another HSA containing fluid, increases growth rate and survival of *A. baumannii* A118 (23).

Changes in the growth of *A. baumannii* A118 were not observed in the presence or absence of CSF in minimal medium containing glucose as a carbon source (Figure S2A). Conversely, the addition of CSF was correlated with an increase in growth rate of *A. baumannii* AB5075 (Figure S2A). The most conspicuous change in expression of an enzyme that participates in the metabolism of glucose was the glucose dehydrogenase, which catalyzes oxidation of glucose to gluconic acid (27). However, expression of this enzyme was increased in *A. baumannii* A118 and decreased in strain AB5075 (log2fold change 0.62, as determined by RNA-seq) when CSF was added to the growth medium. These results do not show a clear correlation of CSF-differentially regulated levels of expression of this enzyme and growth rate. However, an increase in the growth rates of both strains was observed under CSF treatment in minimal medium containing citrate (Figure S2B).

There are numerous reports supporting the hypothesis that increasing the expression of enzymes involved in transcription, translation, and synthesis of ATP is correlated with an increase in bacterial growth rate (28–32). However, these growth differences were not evident in either of the *A. baumannii* strains in the presence of CSF. An attractive hypothesis to explain this observation is that the increase in gene expression capabilities is channeled toward the synthesis of cell components necessary for survival in the human body, e.g., adhesins and pilins.

The data described in this section indicates that certain modifications in the *A. baumannii* metabolism are uncoupled from growth rate. Bacterial cells are characterized by allocating resources to maximize growth according to the needs for each environmental condition (27). When growth curves were performed in nutrient-limiting conditions with metabolites including proline, citrate, and glucose, there was an increase in growth of *A. baumannii* under CSF treatment. This suggests that under a nutrient-depleted condition, *A. baumannii* allocates resources to maximize the efficiency of metabolism instead of maximizing growth. In another study that tested the relationship of gene expression with metabolism and growth, it was observed that when bacterial cells were cultured in poor nutrient medium, there was a higher expression of metabolic proteins, such as enzymes and transporters (27).

Our data also suggests that in depleted medium such as CSF, *A. baumannii* may be allocating all possible resources towards metabolism using an uncoupled metabolism to optimize its survival.

### CSF affects the expression of *A. baumannii* virulence genes

Addition of CSF to *A. baumannii* A118 cultures induces an increase in the expression of a set of genes that code for virulence-associated functions such as type IV pili, iron uptake systems, the type VI secretion system (T6SS), and poly-N-acetylglucosamine (PNAG) production.

Type IV pili participate in microbial adherence as well as motility (gliding or twitching). While *A. baumannii* lacks flagellum-mediated motility, twitching, and surface-associated motility was demonstrated in several strains (28, 29). Numerous studies on twitching and surface-associated motility in *A. baumannii* A118 showed dependence on changes in light and temperature (30) as well as on the components of the growth media. In particular, addition of HSA resulted in increased motility and concomitantly upregulation of the cognate genes (22). Exposure of *A. baumannii* A118 to CSF increased the expression of *pilW* (log2fold 1.22), *pilJ* (log2fold change 0.43), *fimA* (log2fold change 3.09), *fimB* (log2fold change 2.32), and the fimbrial protein precursor AbA118F_3133 (log2fold change 1.91). All of the type IV fimbriae genes have been experimentally shown to be associated with motility, cell adhesion, and biofilm formation (31, 32).

The presence of CSF was also correlated with higher expression of twelve genes associated with the acinetobactin iron uptake system (Figure 2 and Table S1). These genes are part of the ferric-acinetobactin receptor-translocation machinery (*bauABDE, bauA* log_2_fold change 1.67), the acinetobactin biosynthesis (*basBDFGJ, basD* log_2_fold change 1.86) and export (*barB*) (33) (34) (Figure 2A and Table S1). Besides their direct role in iron uptake in the iron starvation conditions found in the human host, the products of *basD* and *basA* are needed for *A. baumannii* to persist and cause apoptosis of human alveolar epithelial cells (35). Bacterial iron uptake systems that are virulence factors are usually highly regulated and are induced in conditions of iron starvation. It is was of interest that in *A. baumannii* A118, genes that code for functions in siderophore biosynthesis, export, and import are upregulated of in the presence of CSF. This finding adds another regulatory signal that enhances expression of acinetobactin iron uptake system. This increase in expression could be directly related to growth in the host or to biofilm formation, which is dependent on efficient iron uptake (36).

**Figure 2.**
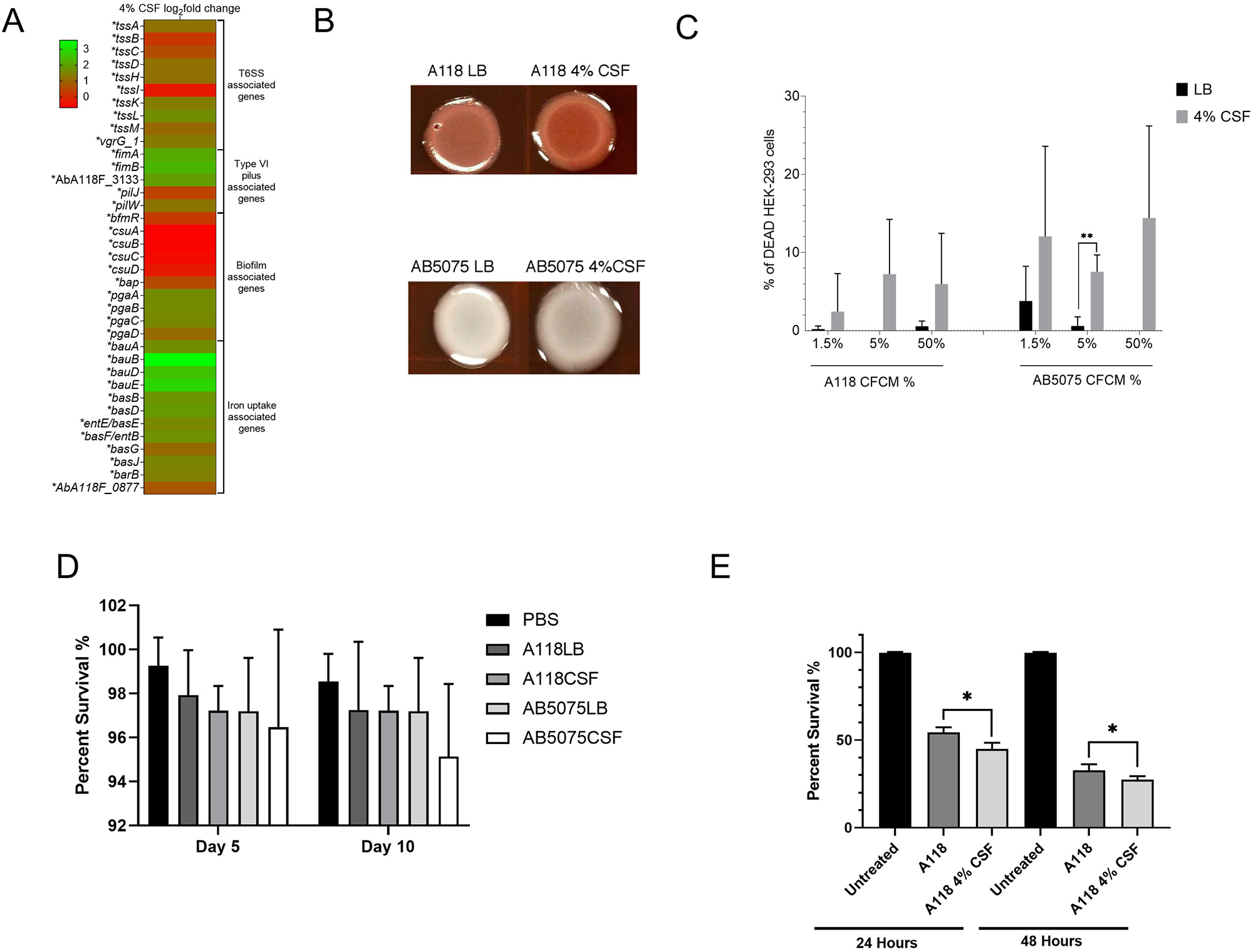
Exposure to CSF can affect multiple virulence factors in *A. baumannii*. A) Heat map of multiple virulence factor associated genes that were differentially expressed in *A. baumannii* strain A118. Asterisks represent a *P-value* of <0.05. B) Poly-N-acetylglucosamine (PNAG) assays were conducted with strains A118 and AB5075 in LB or LB supplemented with 4% CSF. C) Percentage viability of HEK-293 cells under exposure to various concentrations of *A. baumannii* strains A118 or AB5075 supplemented with or without 4% CSF. D) Percentage survival of *Galleria mellonella* when inoculated with *A. baumannii* strain A118 with or without 4% CSF. E) Percentage survival of *Drosophila melanogaster* flies when inoculated with *A. baumannii* strains A118 or AB5075 supplemented with or without 4% CSF.

The T6SS of many bacteria evolved to become essential for virulence (37). They are composed of 14 genes that code for the three components, the phage tail-like, the base plate, and the membrane complexes. Nine out of the 14 *A. baumannii* A118 T6SS genes were slightly but significantly upregulated in the presence of CSF. These genes *tssABCDHIKLM*, code for essential components of the T6SS (Figure 2A and Table S1).

The structures of bacterial biofilms are usually dependent on polysaccharides such as poly-β-1,6-*N*-acetyl-d-glucosamine (PGA) or cellulose. Previous studies showed that functional production of PGA in *Escherichia coli* depends on the products of four genes, *pgaABCD. pgaC* and *pgaD* are essential for biosynthesis, and *pgaB*, which specifies a *N*-deacetylase, together with *pgaA* are needed to export the polysaccharide from the periplasm to the extracellular milieu (38). All four homologs were significantly upregulated when *A. baumannii* A118 was cultured in the presence of CSF (log2fold of 1.63, 1.63, 1.56 and 1.08 for *pgaA, pgaB, pgaC* and *pgaD*, respectively) (Figure 2A and Table S1). As expected, Congo red staining showed that *A. baumannii* A118 cells cultured in the presence of CSF produced higher levels of PGA (Figure 2B).

### CSF enhances the release of *A. baumannii’s* cytotoxic agents

An initial assessment of the effect of CSF on *A. baumannii* virulence was determined using cytotoxicity assays. Cell-free conditioned medium (CFCM) obtained from *A. baumannii* A118 and AB5075 cultured in LB with or without CSF was added to human embryonic kidney cells (HEK-293), and the cells were inspected after 1 hour. Figure 2C shows that CFCM samples obtained from CSF-containing *A. baumannii* AB5075 cultures were significantly more cytotoxic than CFCM from cultures that lacked CSF. This increase in cytotoxicity was observed at all tested concentrations, 1.5% CFCM (*P*-value = 0.006), 5% CFCM (*P*-value = 0.001), and 50% CFCM (*P*-value < 0.0001). Conversely, CFCM obtained from *A. baumannii* A118 cultures containing CSF showed an increased cytotoxic effect only at the highest concentration tested (50%) (*P*-value = 0.002). Although at different levels, the results of these assays suggest that the presence of CSF induces the release of one or more cytotoxic substances by *A. baumannii* (Figure 2C).

### CSF-treatment increases *A. baumannii* virulence

To effect of CSF on *A. baumannii’s* virulence was tested using two models of infection. *Drosophila melanogaster*, recently proposed as a promising model to investigate *A. baumannii’s* interaction with host cells (39), individuals were inoculated with *A. baumannii* A118 or AB5075 cultured in LB medium containing 4% CSF, and inspected at five and ten days post infection. Both strains showed no significant increase in killing after five days but *A. baumannii* AB5075 did show a significant increase in killing after ten days. Survival with respect to individuals inoculated with cells cultured in LB being was reduced by 3.15% (*P*-value=0.186) (Figure 2D). This study was done using a highly robust *D. melanogaster* population previously shown to have higher stress resistance than commonly used inbred fly lines (40).

Infection assays using the *Galleria mellonella* model showed a statistically significant difference in the killing between *A*. *baumannii* A118 cultured in LB or LB plus 4% CSF. As in the *D*. *melanogaster* model, the bacteria cells cultured in the presence of CSF were more virulent (Figure 2E). These results showed a correlation between the transcriptional changes in virulence gene expression observed *in vitro* with an increase in virulence as determined by tests using two different models of infection.

### HSA plays a role in *A. baumannii* pathoadaptation when exposed to CSF

Previous work showed that the presence of PF is correlated with modifications of the expression of more than 1100 *A. baumannii* genes including many virulence factors such as motility, biofilm formation, efflux, T6SS, fibrinolytic activity and capsule genes (18), and with an increase in cytotoxicity and immune evasion (23). HSA, a component of PF, might be responsible for all or part of these effects (18, 20, 22). The experiments showed in previous sections indicate that CSF produces effects similar to those observed with PF such as an increase in cytotoxicity and changes in expression of virulence genes. Both fluids share as major component HSA; other CSF components are glucose (50-80 mg/dl), ions (Na^+^, K^+^, Ca^2+,^ Mg^2+,^ Cl^−^, HCO_3_^−^), low levels of urea, cholesterol, lactic acid, and others (41, 42). An attractive hypothesis is that as it is the case with PF, part of the effects caused by CSF are caused by its HSA content.

To test this hypothesis, *A. baumannii* A118 cells were cultured in LB or LB supplemented with one of the following: CSF, HSA-depleted CSF (dCSF), or HSA. Total RNA was extracted from cells growing in all four conditions, retrotranscribed, and the cDNA was used as template for quantitative polymerase chain reactions (RT-qPCR). We assessed levels of expression of the Type 1 fimbrial protein (*fimA*), *rpmC*, and *atpB* (ATP synthase beta (AbA118F_0480), which were among the most DEGs when CSF was present in the culture broth. The addition of CSF to LB increased transcription levels while addition of dCSF resulted in a reduction in levels of expression of *fimA and rpmC* while little to no changes in expression were observed for *atpB* (Fig 3A, B and C respectively). The enhancing effect observed in the presence CSF was even more pronounced when the cultures took place in LB containing HSA. In this case, *fimA, atpB, and* AbA118F_2933 expression levels were 8-fold, 109-fold, and 6-fold higher, respectively (Figure 3A-C).

**Figure 3.**
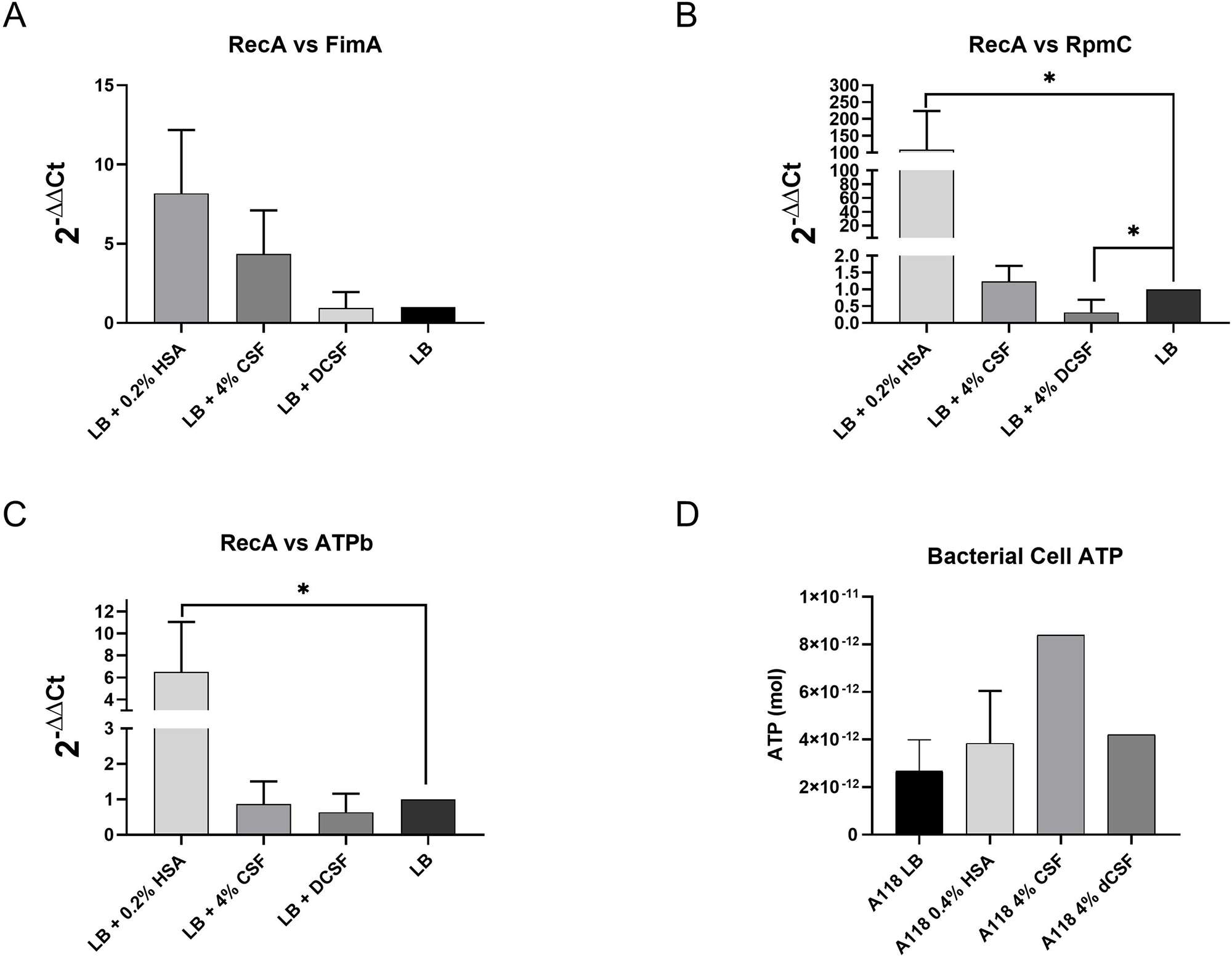
HSA is an essential component for the differential expression of genes in *A. baumannii*. *A. baumannii* strain A118 was exposed to LB, LB plus 0.2% HSA, LB+ 4% CSF and LB + 4% dCSF and its cDNA was synthesized. RT-qPCR was conducted with several genes A) *fimA* encoding gene, B) *rpmC*, and C) *atpB* to determine if HSA was responsible for the increased expression in these overexpressed genes. D) ATP assays were conducted with strain A118 and intracellular ATP was measured to determine if HSA increased the production of bacterial cell ATP.

The expression levels of FoF1-ATP synthase, as determined by ATP synthesis in cells cultured in LB containing CSF or LB containing has, were significantly higher than those observed when the growth media was LB or LB supplemented with dCSF (Figure 3D). These results are in agreement with those obtained by transcriptomics and RT PCR (Table S1, Figure 3C).

Bacterial cells turn on genes that code for factors that allow growth in the hostile environments they encounter upon invading the human body. The results shown in this section indicate that HSA is the signal that triggers the expression of several *A. baumannii* genes when the bacterial cells are in contact with HSA-containing fluids, PF or CSF. Furthermore, our previous studies showed that HSA also enhances DNA acquisition through modulation of natural competence-related gene expression and affects the expression of genes related to motility, efflux pumps, pathogenicity and antibiotic resistance, among others (19, 22). These characteristics are not unique to *A. baumannii*, various bacterial pathogens and protozoa (43–46) modify gene expression to adapt and thrive within the host utilizing HSA as one of the signals. For example, in *Bordetella pertussis*, the causative agent of the whooping cough, albumin combined with calcium induces an increase in production and release of the major toxin, ACT (46). Another example is the case of *Pseudomonas aeruginosa*, in which the presence of albumin is correlated with increased expression of iron-controlled genes (*pvdS* and *regA*) (47). In summary, our observations, together with the evidence available from studies with other bacteria, suggest an important role of HSA as signal for expression of genes and systems essential for survival within the human body.

## CONCLUSION

*A. baumannii* is one of many causative agents of bacterial meningitis, an infection associated with high morbidity and mortality rates. During the infection, bacteria can be found in CSF. In fact, of the several methods available for diagnosis, CSF culture is the most favored (Wu et al BMC 2013 13:26). This study describes changes in expression of numerous genes when *A. baumannii* is exposed to CSF (Figure 5). These genes code for a variety of proteins that participate in the gene expression machinery, energy production, motility, metabolism, survival, and virulence factors among others (Figure 4). Our results in combination with previous work strongly suggest that HSA is a major signaling factor. HSA is one of the main components of CSF and is also present in blood and PF, all body fluids that trigger similar responses in bacteria. Utilizing HSA as the signal for gene expression of elements that facilitate progression of the infection is an intelligent strategy that permits bacteria to sense the presence of human environments. However, not all strains respond equally when HSA is present, slight differences were identified when comparing *A. baumannii* strains A118 and AB5075. These changes are correlated with differences in levels of pathogenicity and probably the kind of infections that are more commonly caused by each variant. An additional remarkable effect of HSA on *A. baumannii* is the augmentation of natural competency (22, 48). Traglia et al. proposed that a random coil stretch in the structure of HSA is responsible for increasing the ability of *A. baumannii* to take up DNA. Considering the pleiotropic effects caused by the presence of HSA on *A. baumannii*, an alternative path to design therapeutic agents against this infection could be to identify compounds that interfere with the ability of *A. baumannii* to detect HSA. Compounds that interact with the HSA regions that are detected by *A. baumannii* could mask the presence of the protein impeding expression of the necessary systems for survival and progression of the infection.

**Figure 4.**
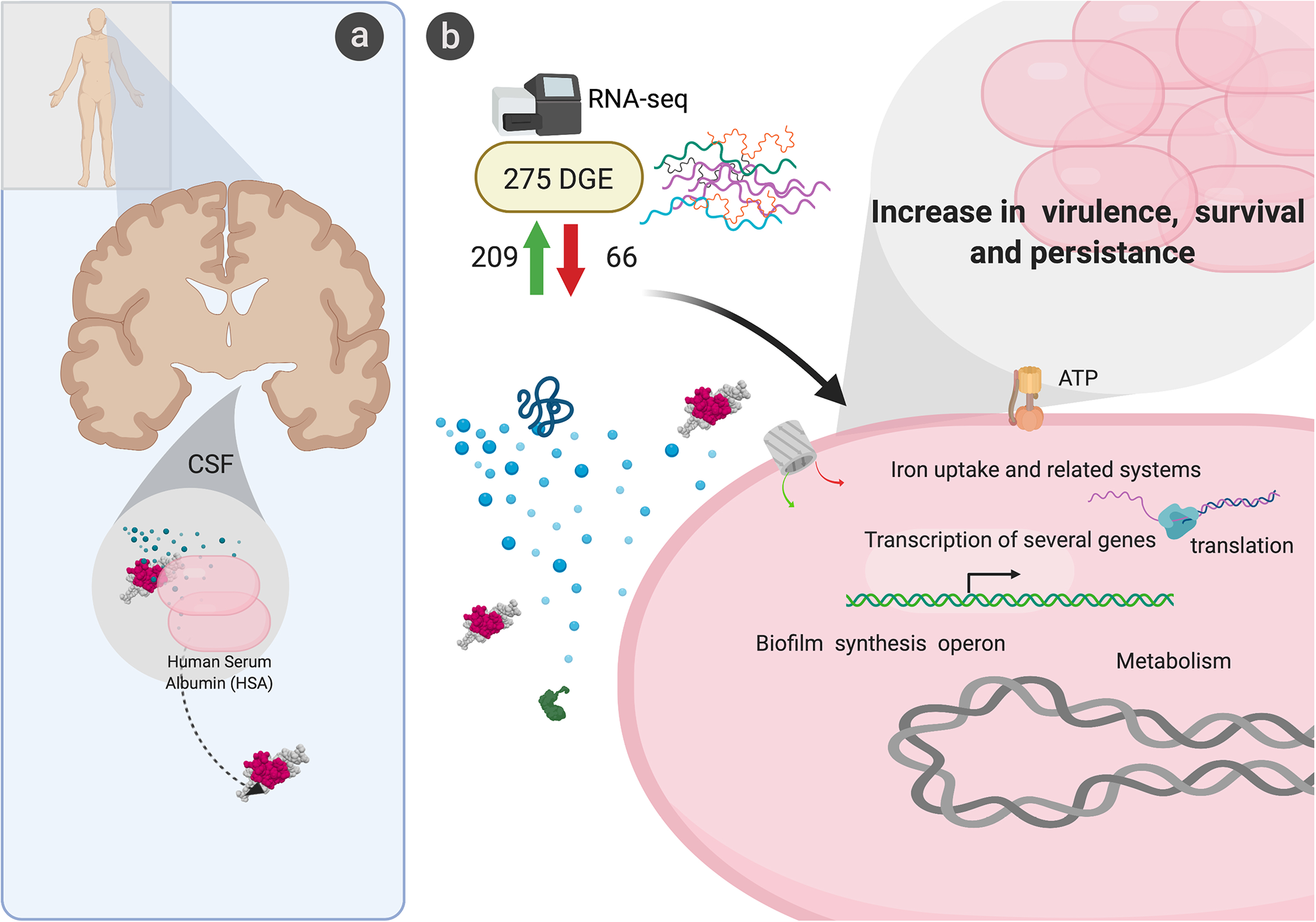
Graphical representation of *A. baumannii’s* transcriptomic and behavioral response to CSF.

## MATERIAL AND METHODS

### Bacterial strains

Two *A. baumannii* strains already used in previous studies (ALL references), exhibiting different degree of susceptibility and virulence were used. A118 strain is known to be susceptible to variety of antibiotics (49, 50) and AB5075 possesses high virulence and is resistant to many antibiotics (51).

### RNA extraction and sequencing

*A. baumannii* colonies (A118 and AB5075) were suspended in Luria Bertani broth (LB) with or without 4% CSF and incubated with agitation for 18 h at 37°C. Overnight cultures were then diluted 1:10 in fresh LB broth and incubated with agitation for 7 h at 37°C. RNA was extracted from each strain using the TRI REAGENT^®^ Kit (Molecular Research Center, Inc., Cincinnati, Ohio, USA) as previously described (22). Total RNA extractions were performed in two biological replicates for each condition.

RNA sequencing was outsourced to Novogene (Novogene Corporation, CA) for mRNA-seq analysis, which includes rRNA depletion, library preparation through Next^®^ Ultra™ RNA Library Prep Kit for Illumina (New England Biolabs) and HiSeq 2500 paired-end 150 bp sequencing.

### RNA-seq Data Analysis

RNA-seq reads (GEO accession No GSE153967) corresponding to *A. baumannii* strain A118 and AB5075 exposed to LB or LB plus 4% CSF were analyzed as follows. Trimming of low-quality bases at the ends of the reads to a minimum length of 100 bp and removal of Illumina adaptor sequences was performed using Trimmomatic (52). FastQC (www.bioinformatics.babraham.ac.uk/projects/fastqc/) was used to assess the quality of the reads before and after trimming. Burrows-Wheeler Alignment software (BWA) was used to align the RNA-seq reads to sequences of the whole genome shotgun sequencing project of strain *Acinetobacter baumannii* A118F (DDBJ/ENA/GenBank accession VCCO01000000). FeatureCounts was used to calculate the read counts per gene, and differential expression analysis was performed using DEseq2 (53, 54). Principal component analysis (PCA) and gene expression heat map with clustering dendrograms of the RNA-seq data analysis of LB and CSF treatments are shown in Supplementary Figure S3. Features exhibiting FDR <0.05 and log2fold change >1 were considered statistically significant.

### Gene ontology analysis

iGO terms were retrieved from UniProt for the best BLASTx hits to *A. baumannii* A118F genes. Using GO.db Bioconductor annotation data package in R language, GO terms and ancestor terms were assigned for all DEGs from this study. GO enrichment analysis was performed using custom-made scripts as described previously (55). The enrichment factor was estimated as the ratio between the proportions of genes associated with a particular GO category present in the dataset under analysis, relative to the proportion of the number of genes in this category in the whole genome. *p*-values were calculated using the Fisher Exact Test and adjusted by the Benjamini-Hochberg method.

### Growth Curves

Both strains, AB5075 and A118, were cultured overnight under different conditions (LB broth and LB broth + 4% CSF). OD_600_ was adjusted to 0.01 using M9 minimal media and used for growth curves with different amino acids at 10 mM (Glutamine and Proline) and carbon sources at 5 mg/mL (Glucose and Citrate). A microplate reader was used to incubate the cultures at 37°C with shaking and OD600 readings were taken every 15 minutes for 24 hours. Growth rate data was then analyzed using Prism 8 software.

### Human Serum Growth Curves

Overnight cultures of A118 and AB5075 cultured in LB broth and LB broth + 4% CSF were OD_600_ adjusted (0.001) in in M9 minimal media and incubated along with either 10% human sera or 10% heat inactive human sera. Human sera were inactivated by placing human sera in a 56°C water bath for 30 minutes. Cultures were incubated at 37 °C with shaking and OD600 readings were taken every 15 minutes for 24 hours. Growth rate data was analyzed using Prism 8 software.

### Cytotoxicity assays

In a Nunclon™ Delta Surface opaque 96-well microplate (ThermoScientific), we added colorless DMEM, 4% CSF, and A118 or AB5075 CFCM diluted in LB broth to make 50 μL of CFCM at final concentrations of 1.5%, 5%, and 50%. An additional 50 μL of ATCC HEK-293 cells at a concentration of 1 × 10^6^ cells/mL in colorless DMEM were suspended in the well and intoxicated for 1 h at 37°C, 5% CO_2_. CellTiter-Glo^®^Reagent (100 μl) was added to each experimental and standard curve well and then placed on an orbital shaker for 2 min. Following mixing, plates were incubated at room temperature for 10 min to stabilize the luminescent signal. The viability of HEK cells was measured at room temperature using the “all” luminescence function of SpectraMax M3.

#### Galleria mellonella infection model

To assess the virulence of *A*. *baumannii* with and without CSF *in vivo*, the *G*. *mellonella* insect model of infection was used (56). Larvae weighing between 200 and 400 mg were maintained on wood chips in the dark at 4°C. *A*. *baumannii* A118 was grown overnight in either LB or LB with 4% CSF. An equivalent of 1.0 OD_600nm_ unit of overnight culture was pelleted and resuspended in 1 mL of cold sterile 20 mM phosphate buffered saline, pH 7.4 (PBS). The cells were further diluted 1:10 in sterile PBS and used for injections. A Hamilton syringe was used to inject 5 μL of the diluted bacterial suspension via the left proleg of each larva. A control group of untreated larvae was used to assess overall larval viability for the duration of the assay. One hundred *G*. *mellonella* larvae were used in each condition and incubated at (37 °C) in a sterile Petri dish for 24 h intervals for 48 h total. Larvae viability was monitored by observing response to gentle prodding with a glass rod; those with no response were considered dead. Four replicates with 100 larvae per Petri dish were performed for each condition.

#### Drosophila melanogaster model of infection

The *D. melanogaster* population used in this study, population B1, is a large outbred population, maintained on 14-day discrete generation cycles, 24:0 light:dark, 25°C, on banana molasses diet (40, 57). Eggs were collected at densities of ~60 eggs/vial. Freshly eclosed adults were transferred to fresh food vials. On day 14 from egg (~4-5 days post eclosion), male flies were inoculated with: PBS, A118 LB, A118 4% CSF, and AB5075 LB, AB5075 4% CSF. All bacterial suspensions were grown overnight and diluted in PBS to an OD of 1. To carry out the inoculation, a stainless steel pin (Fine Science Tools 26002-10, length 1cm, tip diameter 0.0125mm, rod diameter 0.1mm) was sterilized with ethanol, then dipped in each bacterial suspension (or PBS control) and pricked into the ventrolateral thorax of each fruit fly, following the pricking protocol described in Khalil et al. 2015 (58). After inoculation, flies were maintained in groups of ten same-sex flies per vial on banana-molasses medium. Three hours after inoculation, flies that died from handling were discarded, which was always <5% of the flies. Thereafter, flies were transferred to fresh diet every three days. Survival was recorded daily. 50 male flies were inoculated per treatment per each of three biological replicates.

### PNAG assays

To study extracellular matrix (ECM) production, microcolony biofilm was used as model system. 5 ul of overnight cultures of A118 cultured in LB broth and LB broth + 4% CSF were inoculated on LB agar and supplemented with Congo red as previously described (59). Plates were incubated at 28°C in static incubator for up to 48hs. Results were record at 24 hours with a Plugable USB 2.0 Digital Microscope.

### HSA depletion

HSA was depleted from CSF by placing 1mL of CSF into a 30 kDa Amicon™ Ultra Centrifugal Filter (Millipore, Temecula, CA, United States) and the solution was centrifuged at 20,000 × g for 10 minutes. To identify HSA was successfully depleted, an SDS-PAGE was conducted that contained CSF, depleted CSF (dCSF), and pure HSA.

### qPCR

Previously extracted and DNase-treated RNA from *A. baumannii* strain A118 grown in LB, 4% CSF, 4% depleted CSF and 0.2% HSA, were synthesized to cDNA using the manufacturer protocol provided within the iScript™ Reverse Transcription Supermix for qPCR (Bio-Rad, Hercules, CA, United States). The cDNA concentrations were measured with a DeNovix DS-11+ spectrophotometer; each sample was then diluted to a concentration of 50 ng/μl. qPCR was conducted using the iQ™ SYBR^®^Green Supermix through the manufacturer’s instructions. At least three biological replicates of cDNA were used and were run in quadruplet. All samples were then run on the CFX96 Touch™ Real-Time PCR Detection System (Bio-Rad, Hercules, CA, United States).

The transcript levels of each sample were normalized to the *recA* rRNA transcript levels for each cDNA sample. The relative quantification of gene expression was performed using the comparative threshold method 2^−ΔΔCt^. The ratios obtained after normalization were expressed as folds of change compared with cDNA samples isolated from bacteria cultures on LB. Statistical analysis (Mann–Whitney test) was performed using GraphPad Prism (GraphPad software, San Diego, CA, United States). A *P*-value < 0.05 was considered significant.

### ATP assay

Both strains, A118 and AB5075, were cultured overnight in LB broth with or without 4% CSF. Cultures were then diluted 1:100 using fresh LB broth and were incubated at 37 °C with agitation. After 6 hours of incubation, an aliquot of each sample was taken. The OD600 of each sample was recorded. Samples were prepared for the ATP assay using the perchloric acid extraction method as previously described. 100 ul of ice-cold 1.2 M perchloric acid was added to 200 ul of the bacterial sample and they were vortexed for 10 seconds. The mixture was then incubated on ice for 15 minutes. After incubation, the samples were spin down at 16,100 × g for 5 minutes at 4°C. 200 ul of the supernatant was then transferred into a fresh tube and mixed with a neutralizing solution (0.72 M KOH and 0.16 M KHCO_3_). The neutralized extract was spin down and the supernatant was then transferred into a fresh tube and used for the ATP assay.

To measure intracellular ATP content, we followed the manufacturer instructions (Promega, Madison, WI, United States). Briefly, 100 ul of the supernatant was added into an opaque 96 well plate and allowed to stabilize to room temperature. 100 ul of the Bac-Titer™ Glo Reagent was added into each of the samples and the plate was mixed in an orbital shaker and incubated for 5 minutes. A standard curve was included in each plate. Luminescence of the samples was recorded.

## Supplementary material

**Table S1. Differential gene expression analysis of strain A118 under exposure to 4% CSF.** RNA-seq read counts of 2 biological replicates of LB or LB with 4% CSF treated *A baumannii* A118 were analyzed using DEseq software. For each hit, gene ID, average base mean, base mean group A (LB-treated), base mean group B (LB+CSF treated), fold-change, log_2_fold change, *P-value*, Benjamini-Hochberg adjusted *P-value*, and gene description/function are provided.

**Figure S1.** Minimal media growth curves of strain A118 and AB5075 in A) 10mM of glutamine and B) 10mM of proline.

**Figure S2.** Minimal media growth curves of strain A118 and AB5075 in A) 5mg/mL of glucose and B) 5mg/mL of citrate.

**FIGURE S3. Bioinformatic analysis of RNA-seq data from LB- or 4% CSF-treated *A. baumannii* A118**. A) PCA plot of all RNA-seq samples. Biological replicates of the same treatment are indicated by color in the legend. B) Heat map of the expression profiles of the 200 genes displaying the highest variance across samples based on DESeq2 analysis of read count data. Sample-wise (columns) clustering dendrogram is shown.

